# T7Max transcription system

**DOI:** 10.1101/2021.10.17.464727

**Authors:** Christopher Deich, Brock Cash, Wakana Sato, Judee Sharon, Lauren Aufdembrink, Nathaniel J. Gaut, Joseph Heili, Kaitlin Stokes, Aaron E Engelhart, Katarzyna P Adamala

## Abstract

Efficient cell-free protein expression from linear DNA templates has remained a challenge primarily due to template degradation. Here we present a modified T7 RNA polymerase promoter that acts to significantly increase the yields of both transcription and translation within *in vitro* systems. The modified promoter, termed T7Max, recruits standard T7 RNA polymerase, so no protein engineering is needed to take advantage of this method. This technique could be used with any T7 RNA polymerase-based *in vitro* protein expression system. Unlike other methods of limiting linear template degradation, the T7Max promoter increases transcript concentration in a T7 transcription reaction, providing more mRNA for translation.

## Introduction

The T7 promoter for the RNA polymerase of bacteriophage T7 consists of 18 base pairs of sequence (5’ – TAATACGACTCACTATAG – 3’).[1] Previous research identified the relationship between the sequence and transcriptional efficiency, which helped to strengthen the T7 system’s usability.[2–4]

Due to the T7 system’s versatility, the T7 system can be used both *in vivo* and in a cell-free translation system (CFTS). For example, bacterial cell-free translation systems commonly use the T7 RNA polymerase alongside the endogenous sigma 70 system.[5] Furthermore, cell-free translation system platforms derived from hosts other than bacteria are also coupled with the T7 transcription, like plant[6], mammalian[7], and insect[8] *in vitro* translation systems.

We investigated whether translation in a cell free transcription – translation system (TxTl) can be increased by improving the availability of the mRNA. It has been shown that increasing plasmid concentration directly correlates with increased translation yields in bacterial TxTl.[9] We reasoned that increasing the abundance of mRNA, with all other components of the translation system being equal, should result in both an increase of protein abundance and an increased resistance to DNA template degradation by endogenous nucleases in TxTl.

## Results and discussion

Due to the robustness and high popularity of T7 RNA polymerase, there has been a lot of effort in engineering T7 RNA polymerase promoter sequences.[3,10] We began by investigating efficiency of several known T7 promoter variants (**Table 1**). [11,12] We constructed double stranded linear DNA templates coding for the broccoli fluorescent RNA aptamer [13]with each of the tested T7 promoter variants. The templates had no terminators, so all transcriptions were run-off terminated.

**Table 1.**
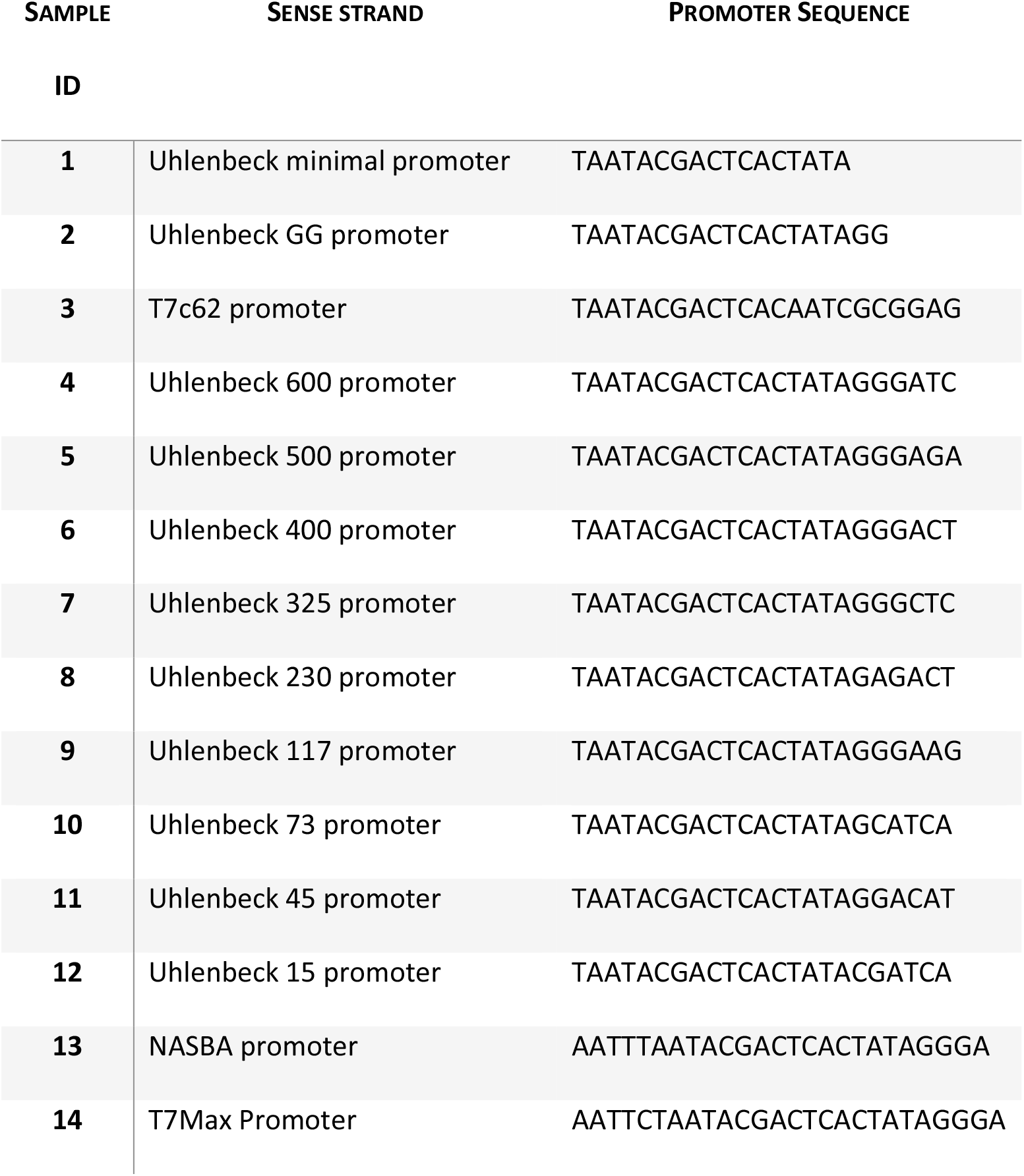

Each transcription reaction was analyzed on a urea PAGE gel with the product stained with DFHBI1T, the ligand for the broccoli aptamer. The resulting image shows only correctly folded full length broccoli aptamer products (**Figure 1a**). The gel was then de-stained and stained again using the general nucleic acid stain Sybr Gold. This stain shows all nucleic acid present in the sample, including truncation products of transcription (**Figure 1c)**. Both DFHBI and Sybr stained gels were quantified, comparing the relative abundance of the full-length broccoli aptamer product to the total nucleic acid abundance in the sample (**Figure 1b** and **1d)**.

**Figure x1:**
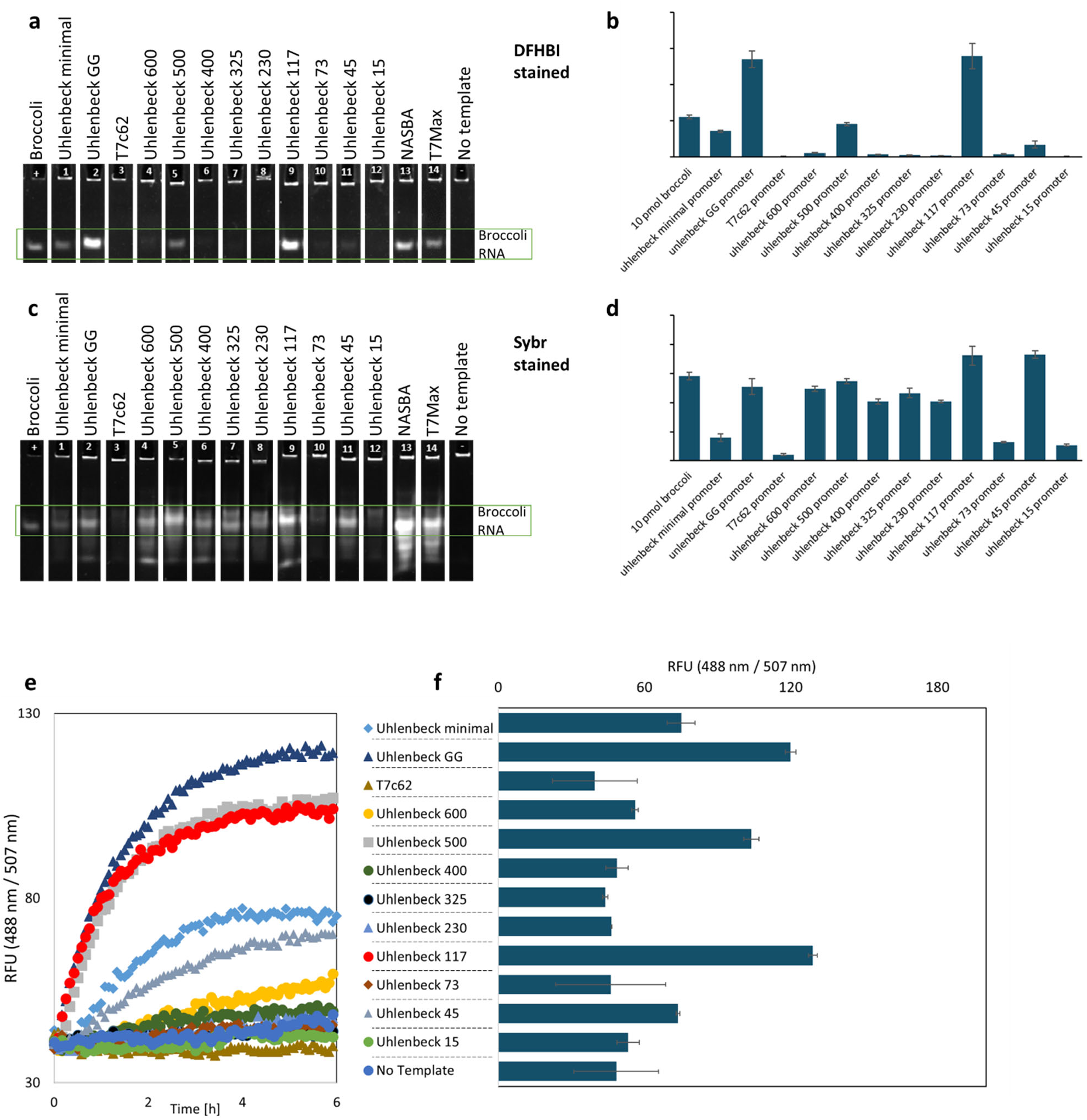
Testing different promoters in *in-vitro* transcription. **a:** transcription of the RNA broccoli aptamer from linear dsDNA templates under different promoters. The gels are stained with DFHBI1T. **b:** quantification of DFHBI1T stained gels. Y axis is the unitless relative brightness of the broccoli RNA band. **c:** quantification of the same transcription gel as in a, stained with Sybr stain. **d:** quantification of the Sybr stained gel. The Y axis is unitless relative brightness of the broccoli RNA band. **e:** time course of transcription from linear dsDNA broccoli aptamer templates with different promoters, one example trace for each experiment. The legend applies to panels e and f. **f:** end point fluorescence of broccoli RNA aptamer for 3 replicates for transcriptions showed on panel **e**, error bars are standard deviation.

We identified two promoters (Uhlenbeck GG and Uhlenbeck 117) that show the highest yields of fluorescent RNA product. We also performed time course fluorescent readout of transcription from all the tested promoters, measuring transcription fluorescence for 6 hours (**Figure 1e**, and end point quantification shown on panel **1f**). We selected the two most promising promoters for further tests: Uhlenbeck GG and Uhlenbeck 117.

Next, we proceeded to test full translation efficiency, still using linear dsDNA template. We constructed eGFP templates with each of the tested promoters, using UTR1 and T500 terminator sequences optimized for bacterial *in vitro* translation.[5] The translation efficiency was measured by fluorescence of eGFP after an 8 hour reaction (**Figure 2a**). We quantified the abundance of eGFP mRNA using RT qPCR (**Figure 2b**). One of the promoters provided a slightly higher protein product amount, and higher mRNA abundance. That promoter, with the sequence AATTCTAATACGACTCACTATAGGGA, which we named “T7Max” – is an improved T7 promoter variant.

**Figure x2:**
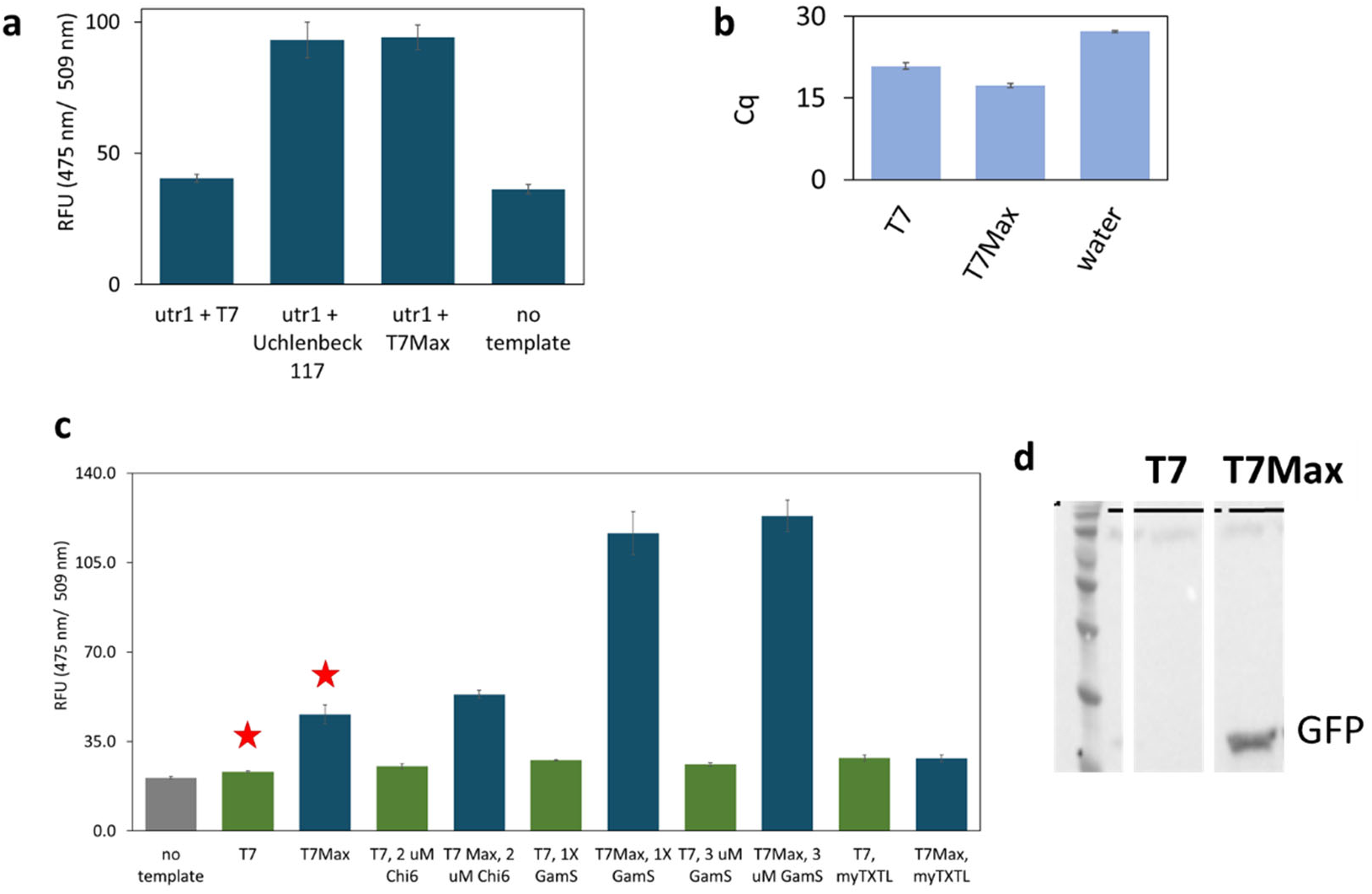
Cell-free TxTl of GFP from dsDNA linear template with different promoters. **a:** cell-free TxTl synthesis of eGFP, with two top candidate promoters, end point fluorescence measured after 8-hour reactions. **b:** RT-qPCR measurement of mRNA abundance in TxTl GFP translation of classic T7 promoter, new T7 Max promoter, and no template control sample. Samples were collected after an 8-hour TxTl reaction. **c:** cell-free TxTl synthesis of GFP, T7 promoter (green bars) and T7Max promoter (blue bars), in house-made bacterial TxTl, with different ways of protecting linear DNA templates, and with commercially available myTXTL kit; end point fluorescence measured after 8-hour reactions. For panels **a, b** and **c:** each sample in triplicate, error bars are S.E.M. **d:** example of Western Blot analysis of GFP translation, 8-hour end point translation from linear dsDNA template in home-made TXTL without DNA protection reagents (samples represent conditions showed on panel c marked with red star).

*Escherichia coli* has many endogenous DNA nucleases[14], which make their way into the TxTl extract without losing activity[15] and thus cause degradation of linear DNA templates in TxTl. Several methods have been proposed for enabling linear template expression, mainly focused on blocking the activity of the RecBCD, one of the more well-characterized nucleases. Among those methods, the most popular are the addition of GamS protein[16] or small DNA Chi6 [17] – both inhibiting RecBCD nuclease.

We used both the Chi6 inhibitors, and the GamS protein inhibitor. We tested expression of eGFP under classic T7 and under T7Max, from the same linear templates described above, using either *E. coli* extract made in our lab, or MyTXTL, a commercial *E. coli* TxTl extract from Arbor Biosciences (**Figure 2c**). All reactions were set up with identical DNA template concentrations and in each compared pair (T7 vs T7Max) all other conditions, like concentration of RecBCD inhibitor, were the same. In all cases, the T7Max promoter outperformed the classic T7 promoter, as measured by GFP fluorescence after an 8 hour reaction. In some cases, expression under the T7Max promoter was 5 times larger than expression under the classic T7 promoter (in cases of GamS experiments, **Figure 2c**). In addition to fluorescence measurements, we confirmed via a Western Blot one sample for each of the tested conditions (**Figure 2d)**.

After establishing that the T7Max promoter outperforms the classic T7 promoter in expression from linear DNA templates, we moved on to further characterizing the T7Max promoter in translation reactions.

We used two circular DNA plasmids using UTR1 and T500 terminator and eGFP, identical except for the sequence of the promoter. First, we compared the kinetics of eGFP translation in *E. c oli* TxTl (**Figure 3a**), and corresponding GFP mRNA abundance (**Figure 3c**).

**Figure x3:**
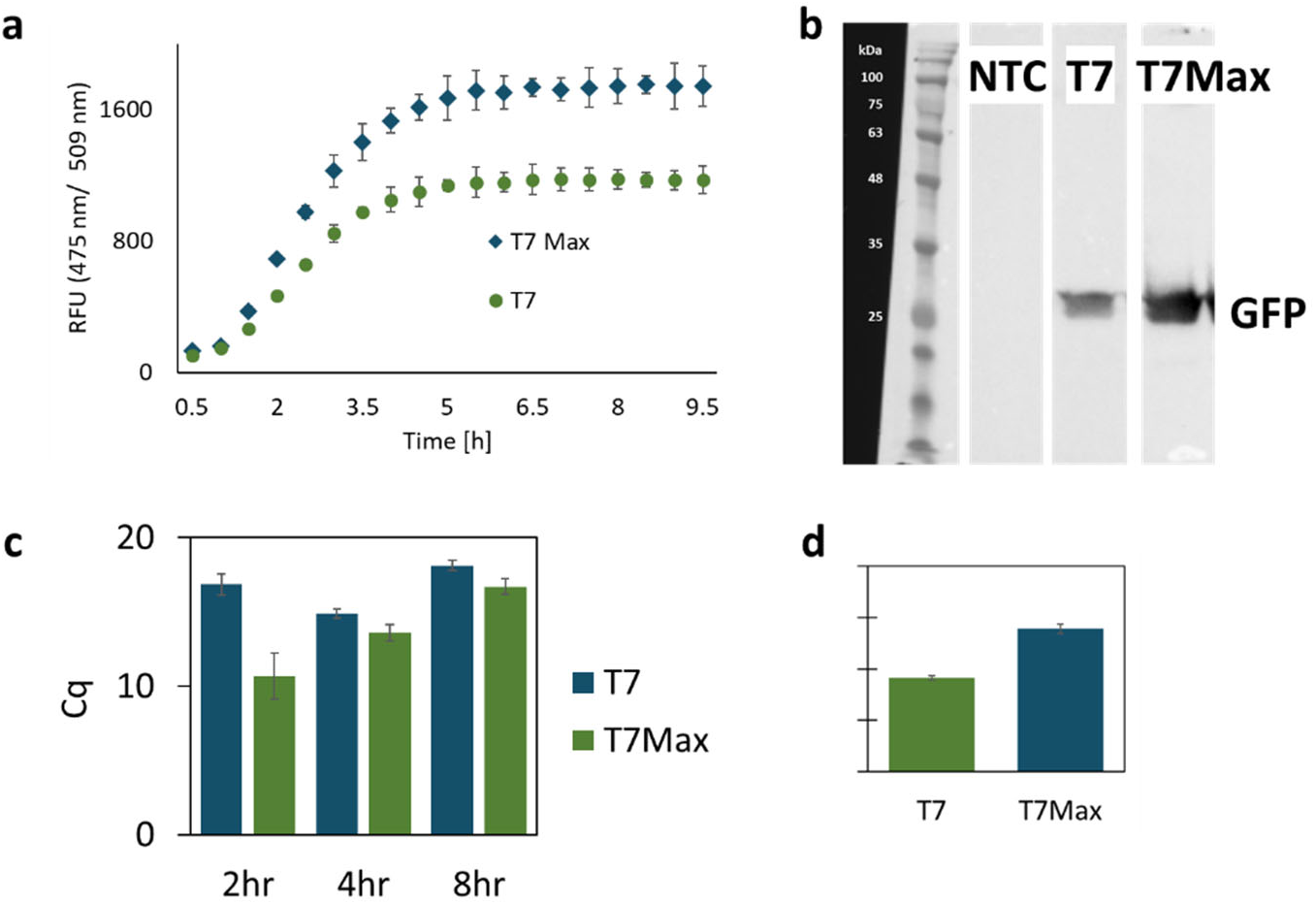
Cell-free TxTl of GFP from dsDNA circular plasmid template with different promoters. **a:** time course expression of GFP under the classic T7 vs T7Max promoter. **b:** Western Blot analysis of expression of GFP. **c:** RT qPCR cycle (Cq) value quantifying abundance of GFP mRNA. **d:** quantification of Western Blots of GFP expression, expressed as unitless relative brightness value. All samples in triplicate, error bars represent S.E.M. Protein product was measured by end point measurements after an 8-hour reaction.

The T7Max promoter consistently provided a higher level of fluorescence and a higher copy number of mRNA than the classic T7 promoter. To ensure that the measured protein abundance is not a fluorescence artifact, we analyzed eGFP abundance via Western Blot (**Figure 3b**) and then quantified the Western Blot gels (**Figure 3d**). The T7Max promoter consistently produced higher protein abundance.

To further characterize performance of the T7Max promoter in cell-free protein expression reactions, we analyzed reactions at different temperatures. In addition to 30°C (the optimal *E. c oli* TxTl reaction temperature used throughout this paper), we analyzed reactions at 25°C and 37°C (**Figure 4a**).

In all cases, T7Max produced more protein product, confirmed by RT qPCR measurements of mRNA abundance (**Figure 4b**). The advantage of T7Max was largest at 30°C, the optimal TxTl temperature, however the qPCR data shows significantly higher abundance of mRNA produced from T7Max vs classic T7 at 37°C as well. We speculate that this discrepancy might be due to the generally decreased translation performance at higher temperatures.

**Figure x4:**
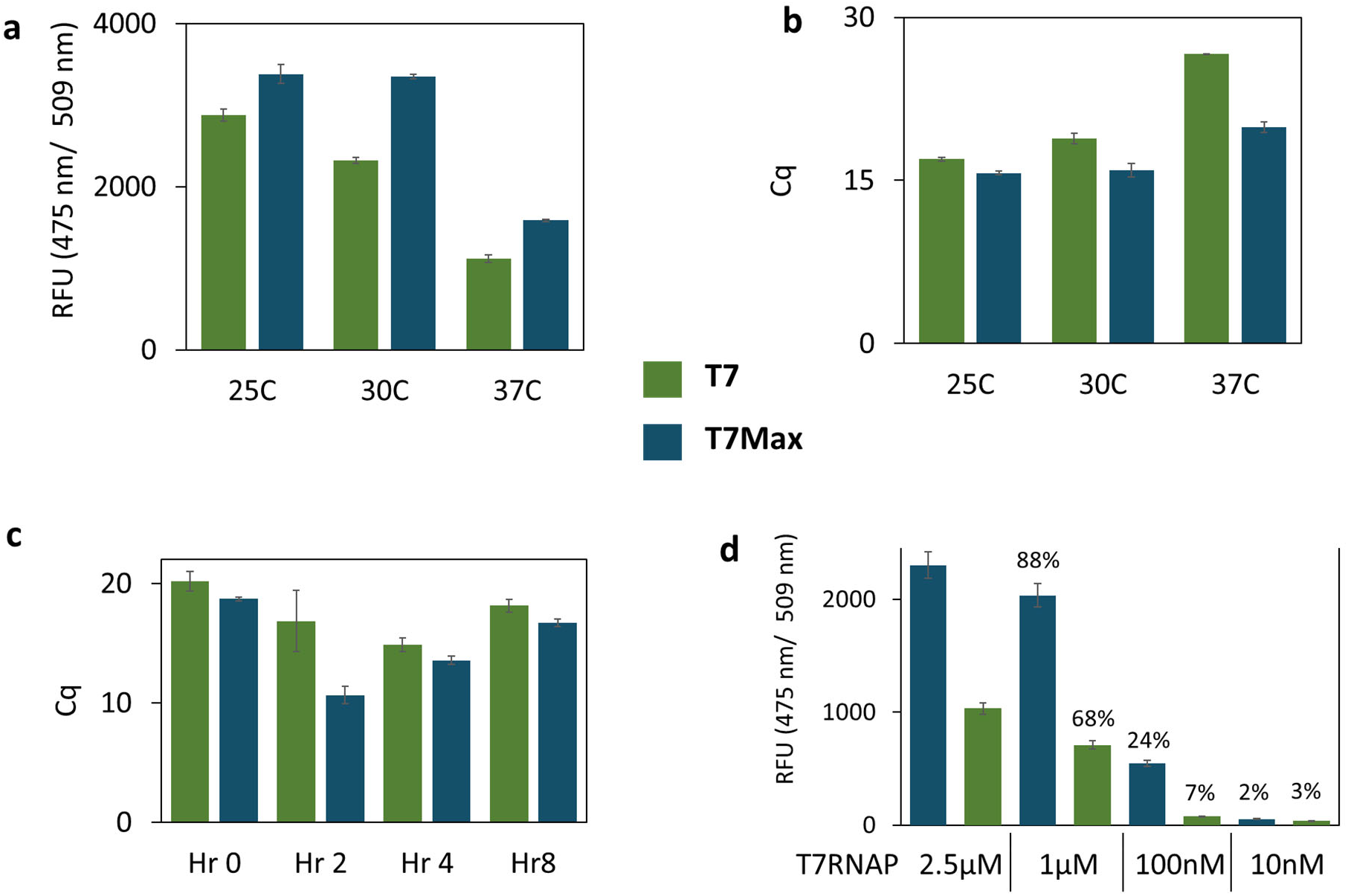
T7Max performance characterization. Translation of GFP protein from circular plasmid template was measured at different temperatures and with different T7 RNA polymerase concentration. All green bars: T7 promoter, all blue bars: T7Max promoter. **a:** expression of GFP measured after an 8-hour reaction at different temperatures. **b:** RT qPCR measuring abundance of GFP mRNA in samples from panel **a. c:** mRNA abundance measured at different times during the TxTl reaction at 30°C. **d:** expression of GFP measured after an 8 hour reaction with different concentration of T7 RNA polymerase. The percentage numbers above bars show fluorescence relative to the value at 2.5μ M T7 RNAP for each promoter. All samples in triplicate, error bars represent S.E.M.

The analysis of mRNA abundance in a TxTl reaction over time (**Figure 4c**) demonstrates that T7Max consistently produces more mRNA than classic T7, however the biggest difference is visible at the 2-hour mark . We speculate this might be due to the interplay between mRNA synthesis and degradation: the mRNA produced from the T7Max promoter is identical to the one produced from the classic T7 promoter, therefore after significantly higher accumulation initially due to faster transcription, the abundance evens out due to similar levels of mRNA degradation.

We also investigated the influence of the T7 RNA polymerase concentration on translation performance (**Figure 4d**). Comparing the T7Max promoter with the classic T7 promoter demonstrates that the T7Max promoter produces higher protein yield at higher T7 RNA polymerase concentrations. However, as the T7 RNA polymerase concentration decreases , the difference between the T7Max and classic T7 templates starts to even out. We speculate this is because at lower RNA polymerase concentrations, the polymerase concentration becomes the rate limiting factor. While T7Max provides more efficient translation, if there is not enough polymerase to bind to all DNA templates, the promoter strength becomes less significant.

To thoroughly characterize the difference in T7Max performance vs classic T7 performance, we expressed several different types of proteins differing in open reading frame size from 1650bp to 30bp (**Figure 5**).

**Figure x5:**
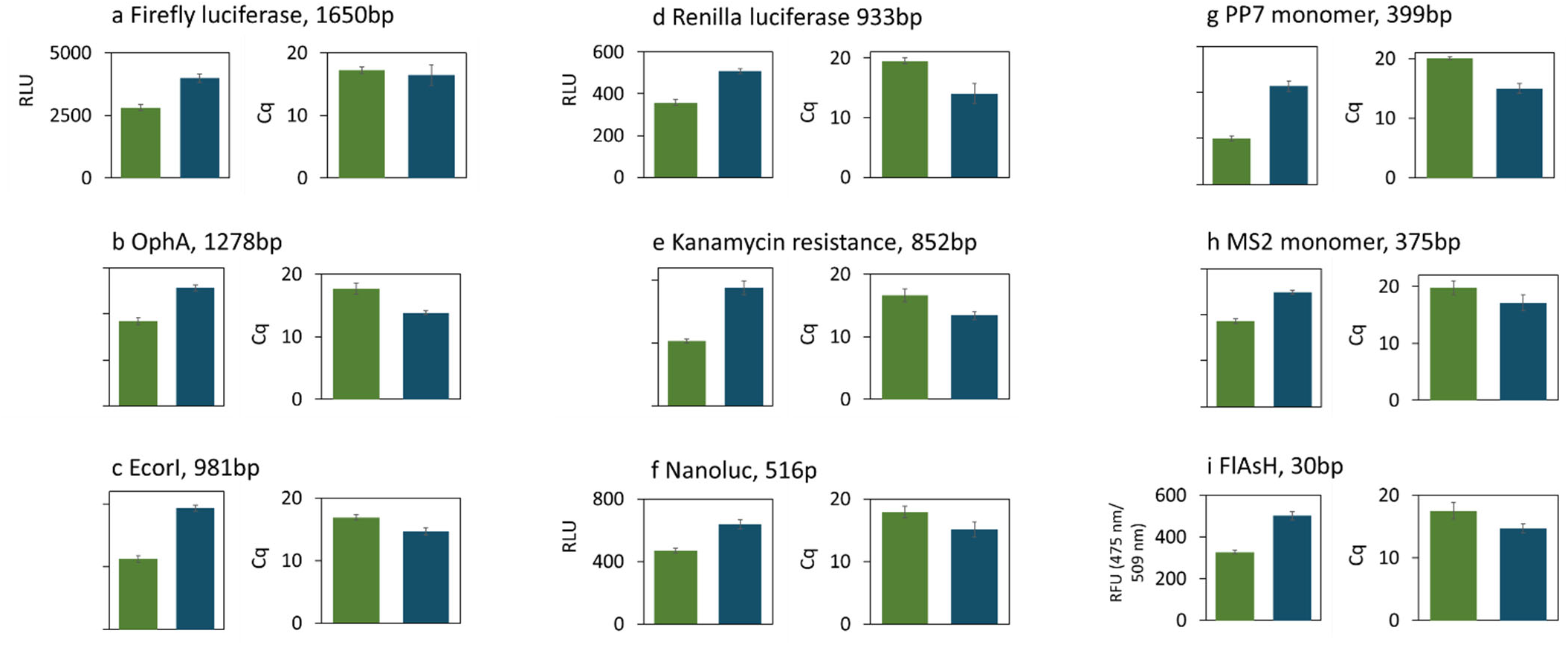
Performance of T7Max vs T7 promoter in different template lengths. All green bars: T7 promoter, all blue bars: T7Max promoter. Circular plasmid DNA template expression of proteins with different length of the open reading frame, from 1650 base pairs to 30 base pairs. Each graph shows protein product quantification and corresponding RT qPCR cycle (Cq) value quantifying abundance of mRNA for each protein. All samples in triplicate, error bars represent S.E.M. Protein product was measured by end point measurements after an 8-hour reaction. Luminescence with appropriate luciferase product was used on panels **a, d, f**. Quantification of appropriate size Western Blot band, expressed as unitless relative brightness value, was used on panels **b, e, g** and **h**. Fluorescence with the arsenic ligand was measured on panel **i**.

We expressed luciferases: firefly (**Fig. 5a**), Renilla (**Fig. 5d**) and Nanoluc[18] (**Fig. 5f**). We expressed viral coat protein RNA binding proteins PP7[19] (**Fig. 5g**) and MS2[20] (**Fig. 5h**). We expressed the protein OphA from *Omphalotus olearius* Jack-o’-Lantern mushroom (**Fig. 5b**). We expressed the DNA restriction enzyme EcoRI (**Fig. 5c**), and the kanamycin resistance protein (**Fig. 5e**). We also expressed the extremely small fluorescent protein aptamer, FlAsH aptamer, which binds an arsenic ligand[21] (**Fig. 5i**). Thus, we covered a wide range of protein sizes, and many possible mRNA folds.

In all cases, in addition to measuring the protein abundance after an 8 hour TxTl reaction, we performed RT qPCR analysis of mRNA abundance. In all cases, T7Max templates produced more protein and higher mRNA abundance than classic T7 templates.

Cell-free translation systems are key components of most synthetic minimal cell designs.[23] We tested the T7Max promoter in the cytoplasm of a synthetic cell : encapsulating *E. c oli* TxTl in POPC / cholesterol liposomes.[22] We prepared samples of synthetic cells with phospholipid membranes, dyed red with Rhodamine-PE dye, and bacterial TxTl with eGFP-encoding plasmid under the control of either the classic T7 promoter or our T7Max promoter. (**Figure 6**). Imaging of the diluted samples clearly showed individual synthetic cell liposomes expressing GFP in the lumen (**Figure 6a and 6b**). To increase the number of samples analyzed in each field of view, we also imaged undiluted samples, at higher concentrations of lipids (**Figure 6c and 6d**). We quantified fluorescence from these images, measuring total fluorescence in the GFP channel to estimate protein production and then normalizing that value to total fluorescence in the red channel (normalizing to the number of liposomes in each field of view). Synthetic cells expressing GFP under the T7Max promoter showed higher protein production than synthetic cells containing the classic T7 promoted GFP.

**Figure x6:**
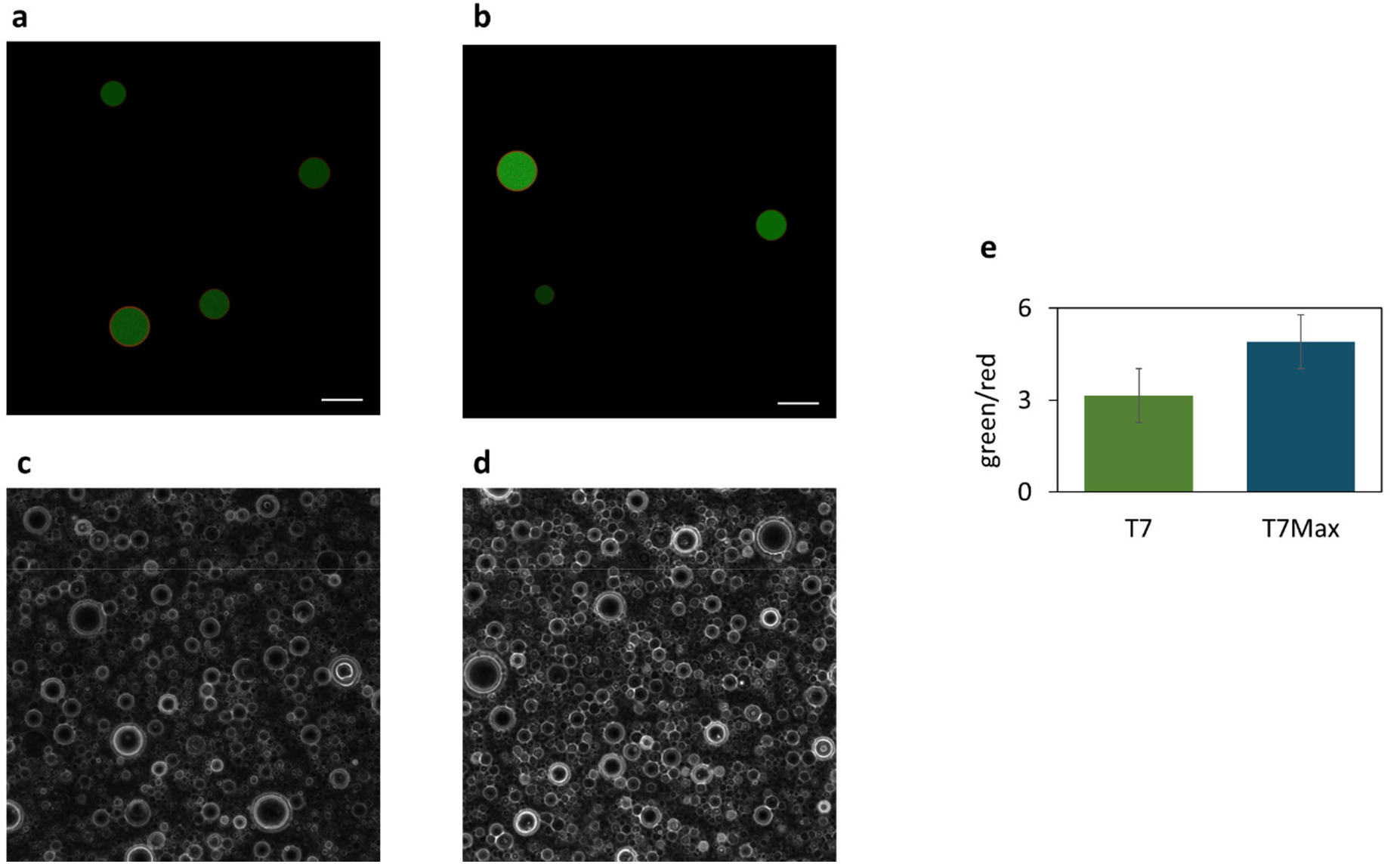
Synthetic minimal cells expressing GFP protein. Microscope images showing liposomes encapsulating plasmid encoding GFP under T7 (panels a and c) and T7Max (panels b and d) promoters. Panels **a** and **b**: 0.1mM lipid concentration, green (GFP) and red (rhodamine membrane dye) channels overlayed. Panels **c** and **d**: bright field showing density of liposomes at 10mM lipid. Scale bar is 5μ m. **e**: quantification of 5 images taken from different fields of view in samples at 10mM lipid; the value is ratio of total fluorescence in green channel to total fluorescence in red channel. Error bars represent S.E.M.

Next, we asked how will T7Max compare to classic T7 in other *in vitro* translation systems. Other *in vitro* translation systems are used for different applications[23,24], including the PURE system composed of *E. coli* translation machinery purified individually[25], wheat germ extract[26], *Leishmania tarentolae* extract[27], insect *Spodoptera frugiperda* Sf21 cell line extract[28], and rabbit reticulocyte extract[29]. All of those extracts are commercially available and were used according to the manufacturer’s protocols.

We created templates for eGFP expression in each of those cell-free systems, with the only difference between templates being the T7 RNA polymerase promoter: either T7Max or classic T7. Because the absolute yields (measured as GFP fluorescence) were different in each extract, we normalized the results: the classic T7 promoter is assigned value 100, and the T7Max template fluorescence is proportionally scaled for each sample. For example, the raw fluorescence value for classic T7 promoter *E. coli* in this case was 9384, while T7Max value was 15671; normalizing T7 to 100 gives T7Max value of 167 (**Figure 7a**). In all tested cases, the yield of protein synthesis was higher from a template using the T7Max promoter than from the template using the classic T7 promoter. Finally, we looked to other applications for T7Max. Robust, sensitive, and transportable disease detection systems are in great need, and many rely on the amplification of nucleic acids.[30] Apta-NASBA is an isothermal exponential disease detection reaction, dependent on the productivity of T7 RNA polymerase.[31] In Apta-NASBA, primers introduce the T7 RNA polymerase promoter and result in a fluorescent read out via an RNA aptamer. We designed primers to detect the *aggR* gene of *E. Coli* – one incorporating T7Max and the other, classic T7. All other reaction components were kept identical. Reactions where T7Max was incorporated created a 14X signal compared to a negative control (a reaction lacking template) vs 1.24X when incorporating classic T7 after 100 minutes (**Figure7b)**. Such increase in signal can allow for a more sensitive detection reaction.

**Figure x7:**
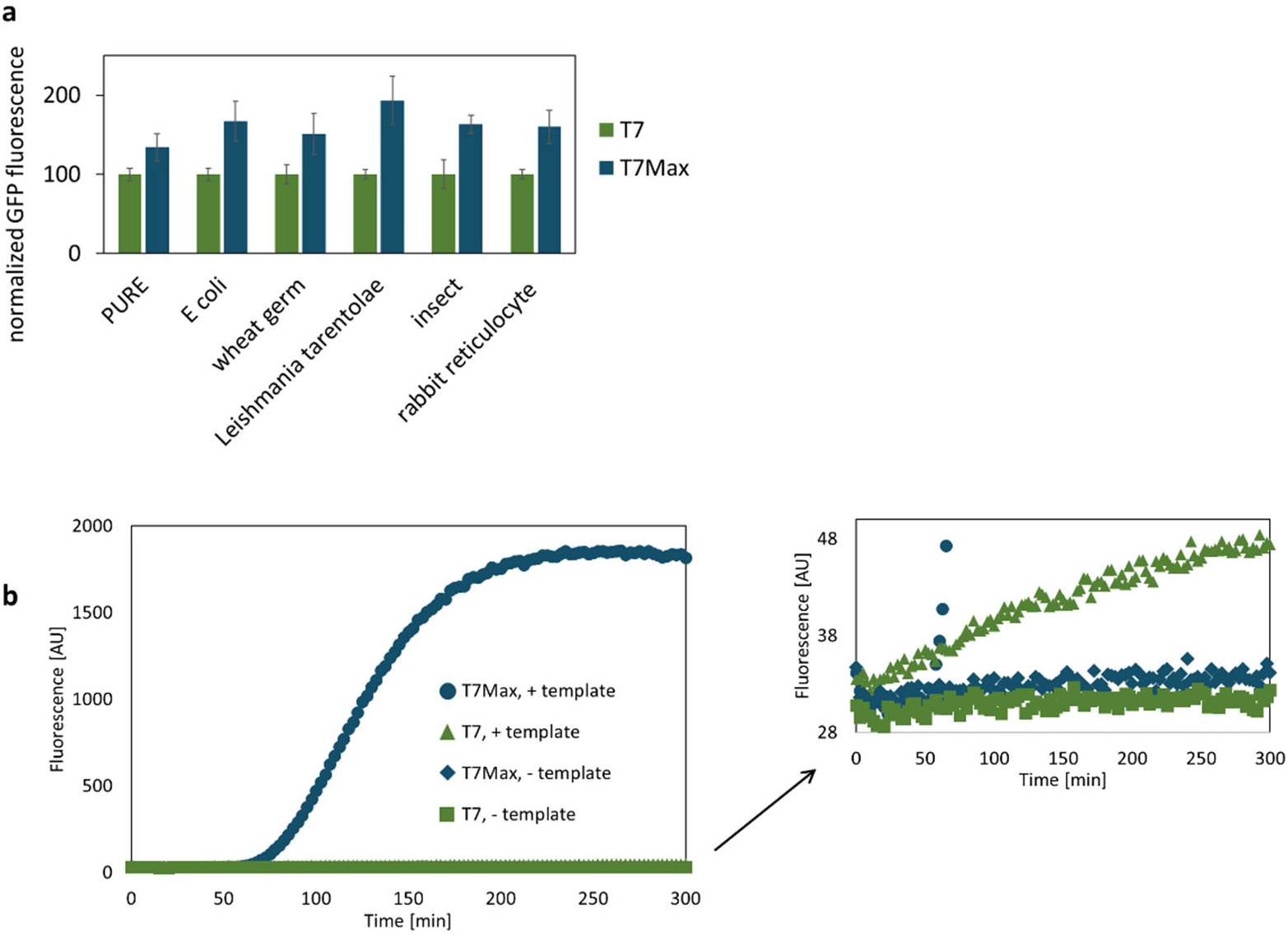
Performance of T7Max in different systems. **a**: Cell-free translation reactions based on different organisms. GFP plasmids were prepared for each specific commercial cell-free expression system (except E. Coli, which used the same plasmids as tested earlier, and in house made cell-free expression system). Fluorescence of GFP protein was measured after each reaction, and raw fluorescence was normalized so that classic T7 promoter fluorescence was assigned value 100, and T7Max sample fluorescence was scaled proportionally. All samples are in triplicate, error bars represent S.E.M. **b**: **b**: Apta-Nucleic Acid Sequence Based Amplification reaction detecting E. Coli gene, aggR. Reactions are identical except for the incorporation of T7Max vs classic T7 promoter. Fluorescence of the broccoli aptamer was measured every 2.5 minutes, excitation: 472 nm and emission: 507 nm. All samples were performed in triplicate, and traces represent the average.

Cell free expression platforms find increasingly versatile applications in many areas of bioengineering, synthetic biology, and metabolic engineering. [32–34] Additionally, the focus for engineering synthetic minimal cells is on reconstituting *in vitro* translation reactions , most often with the use of a bacterial translation system and T7 RNA polymerase.[35,36] Here we demonstrated a simple technique to enable a significant increase in translation yield via a change of the T7 promoter sequence.

This system utilizes all existing elements of T7 RNA polymerase-driven transcription without changes and only requires replacement of the promoter sequence in the construct.

We have demonstrated versatile utility of the T7Max promoter in multiple different cell-free protein expression systems and for proteins over a wide range of sizes and types, as well as significantly increased yields of protein synthesis from linear DNA templates.

With the production of mRNA vaccines, there has been a recent increase of significance for *in vitro* transcription, in particular with the use of the T7 RNA polymerase.[37,38] This increases the need for optimizing transcription reactions. While the sequence of the T7Max promoter has been known for decades, this is the first comprehensive characterization of its use for *in vitro* transcription and translation. Our hope is this technology will enable further improvements in both transcription and *in vitro* protein expression to result in better biomedical, biotechnological and synthetic cell engineering applications.

## Materials and methods

### Construction of Tx Templates for Screen of Different Promoters

A series of T7 promoters described previously [1,2,11,12], see **Table 1** for sequences, were placed upstream of the broccoli coding sequence via primer extension. Sense strand primers with promoter sequences, the first 23 nucleotides of the broccoli coding sequence, and the anti-sense primers coding for broccoli (49 nucleotides) were synthesized (Integrated DNA Technologies) and brought to 10 μ M in Millipore water (GenPure Pro UV-TOC/UF). Bulldog Bio BioReady Taq DNA Polymerase (BSA12L010) was used according to manufacturer’s instructions with NEB dNTPs (N0446S). 9μL of PCR master mix reagents and anti-sense primer were combined with 1 μL of the promoter primer to give a primer-extension reaction of 1X Bulldog Reaction Buffer, 1 μM of both primers, 1 mM dNTPs, BioReady rTaq (0.05 U/μL). The reaction was denatured for 5 seconds at 95°C, annealed for 5 minutes at 60°C, and then extended for 30 min at 72°C (Bio-Rad T100 thermocycler). These reactions were generated in triplicate for each promoter tested, which served as a 10X stock of template in a transcription reaction.

### Transcription

The templates were then used as-is in a transcription assay. All reagents, tubes, and plates were pre-chilled on ice. A master mix of transcription reagents was prepared on ice, and 9 μL of the master mix and 1 μL of the 10X templates were combined in a 200 μ L PCR tube, flicked, spun down, and then transferred to a cold, clear bottom 384-well plate. The transcriptions (1X template, 1X Homemade NEB Buffer, 8 mM GTP, 4 mM A/C/UTP, 0.005X phosphatase 25 ng/μ L, 1 μ M T7 RNAP, 100 μ M DFHBI-1T, RNAse inhibitor 0.4 U/μ L) were incubated for 6 hours at 37°C in a SpectraMax Gemini XS microplate fluorimeter and data collected every 5 minutes (excitation: 472 nm, emission: 507 nm). An endpoint measurement was taken and the transcriptions stored at -80°C.

The fluorescent data was correlated by resolving the transcriptions in a denaturing polyacrylamide gel. An 8M urea, 10% (19:1) PAGE was prerun for 30 minutes at 100V in a Mini PROTEAN tank (Bio-Rad) electrophoresis chamber using 1X TBE (89 mM Tris, 89 mM boric acid, 2 mM EDTA, pH 8.0). Transcriptions were diluted 1:1 with 2x TBE Loading Buffer (8 M urea, 89 mM Tris, 89 mM boric acid, 2 mM EDTA, pH 8.0) and the entire 20 μL sample was resolved for 1 hour at 125V. The gel was then equilibrated in 50 mL 1X folding buffer (1 mM MgCl2, 50 mM KCl, 10 mM Tris, pH 8.0) for 45 minutes. The buffer was then decanted, exchanged with 50 mL 1X folding buffer supplemented with 10 μ M DFHBI-1T, and incubated for 15 minutes at room temperature . The broccoli band was imaged on an Aplegen Omega Lum G using a SYBR Safe filter. The buffer was decanted as before and replaced with 1X Folding Buffer supplemented with 1X SYBR Gold (Thermo Scientific, S11494). After a 15 minute incubation at room temperature , the total RNA was imaged using the aforementioned filter. Low range ssRNA Ladder (New England BioLabs, Cat no N0364S) and 10 pmol of broccoli were run alongside the transcriptions as controls. The RNA produced for both stain s was quantified using GelQuant.NET.

### Construction of T7Max Plasmids

Double stranded T7Max promoter insert was formed from a pair of annealed 5’-phosphorylated primers. Primers were designed with 4 bp 5’ overhangs just upstream of a restriction enzyme digestion site, the forward primer containing the AgeI restriction site and the reverse primer containing the BglII restriction site, using Geneious 7.1.9 (https://www.geneious.com/) and purchased from IDT. For the promoter insert primers, the forward primer sequence was 5’-/5Phos/GATCTAATTCTAATACGACTCACTATAGGGAAATAATTTTGTTTAACTTTAAGAA-3’ and the reverse primer sequence was 5’-/5Phos/CCGGTATATCTCCTTCTTAAAGTTAAACAAAATTATTTCCCTATAGTGAGTCGTA-3’. The T7 promoter sequence was excised from the original plasmid backbone, UTR1-T7RNAP-T500 (Catalog No. 67739, Addgene), via restriction digestion with AgeI and BglII. The T7Max promoter was cloned into backbones containing the genes for eGFP, fluorescein arsenical hairpin (FlAsH) peptide, and Omphalotin A (OphA) by following NEB’s restriction digest protocol (NEB #R0744), 5’ dephosphorylation protocol (NEB #M0289) and T4 DNA ligase protocol (NEB #M0202). Ligated constructs were transformed into the *E. c oli* strain BL21(DE3) and plated on LB agar plates containing 100 μg/ml carbenicillin. Colony constructs were verified by sequencing.

### Western Blot

C-terminus 6xHis -tagged proteins were expressed *in v itro* with transcription-translationally active *E. c oli* cell-free extract using the protocol described before[39]. Constructs were expressed for 8 hours at 30°C using a Bio-Rad T100 thermo cycler running software version 1.201. Samples were mixed 1:1 with 2X SDS loading buffer (100 mM Tris HCl, 2.5% SDS, 20% Glycerol, 4% Beta -mercaptoethanol, 0.1% Bromophenol Blue). Mixtures of loading buffer and sample were boiled at 95°C for 5 minutes in a Bio-Rad T100 thermo cycler. Boiled samples were fractionated on a 37.5:1 Acrylamide:Bis-Acrylamide SDS-Page gel and then transferred to a 0.2 μ m nitrocellulose membrane using a Mini-PROTEAN tank (Bio-Rad) according to the manufacturer’s protocol. Gels were run for 60 minutes at 100V in 800 mL of 1X SDS running buffer (25mM Tris, 192mM Glycine, 3.5mM SDS). Gels were transferred for 60 minutes at 100V in 1L of 1X transfer buffer (25mM Tris, 192mM Glycine). Electrical current was provided by Bio-Rad Power Pac 3000. Membrane was incubated with 5% nonfat milk in TBST (20mM Tris, pH 7.4, 150mM NaCl, 0.05% tween) for 60 minutes on a horizontal rocker (Benchmark) before mouse IgG1 anti-his primary antibodies (1:5000), purchased from Biolegend, were added to the solution. The 5% nonfat milk TBST and mouse IgG1 mixture incubated with the membrane for 60 minutes on a horizontal rocker. After incubation with primary antibodies, the membrane was rinsed three times with TBST followed by three 10 min washes in TBST. The membrane was next added to 5% nonfat milk in TBST containing horseradish peroxidase-conjugate goat anti-mouse IgG1 secondary antibodies (Biolegend 405306) diluted at 1:5000 and incubated on a horizontal rocker for 60 minutes. After incubation with secondary antibodies, the membrane was rinsed three times with TBST followed by three 10 minute washes in TBST. Blots were developed with SuperSignal (Thermo Scientific) immunoblotting detection system according to manufacturer’s protocols. Blots were imaged using the ChemiDoc MP Imaging System (Bio-Rad) running Image Lab version 5.2.1.

### Measuring promoter-dependent protein expression using cell-free TXTL

To prepare the *E. coli* cell extract and TXTL master mix, we followed the protocol outlined by Sun *et al*.[39]. The eGFP, fluorescein arsenical hairpin (FlAsH) peptide, or Omphalotin A (OphA) genes with C-terminal His-tags were cloned into the UTR1-T7RNAP-T500 plasmid backbone (Catalog No. 67739, Addgene). The T7 Max promoter was further cloned into these plasmids for downstream experiments. The linear version of the eGFP plasmid was created through restriction enzyme digestion of the circular plasmid with BamHI. To measure the differences in protein expression between the two promoters, 10nM of templates with each promoter type were added to TXTL reactions and incubated at 30 °C for 8 hours (T100 Thermal Cycler, Bio-Rad). Post-incubation, protein expression was determined through measurement of fluorescence (eGFP and FlAsH) or Western Blot (OphA). eGFP fluorescence was standardized to 1μ M fluorescein.

FlAsH peptide expression was determined through the addition of 5 μ M FlAsH dye and 20mM 2-(*N*-morpholino)ethanesulfonic acid (MES) buffer and were standardized to samples without the peptide. The excitation and emission spectra of FlAsH intersects with that of Chai Green Dye 20X (Catalog No. R01200, Chai Bio) in the subsequent quantitative polymerase chain reaction experiments, so 10 μL of the peptide’s TXTL reactions were saved for transcript quantification prior to determining expression levels.

### Relative comparison of transcripts with Reverse Transcription-quantitative Polymerase Chain Reaction (RT-qPCR)

Template DNA in 10 μ L of the TXTL reaction was degraded by adding 0.5 μ L of TURBO DNase (2U/μ L, Catalog No. AM2238, Invitrogen). The mixture was incubated at 37°C for 30 minutes. The enzyme and the expressed proteins were inactivated by adding 15mM EDTA (Catalog No. E9884, Sigma-Aldrich) at 75°C for 10 minutes (T100 Thermal Cycler, Bio-Rad). The denatured proteins were pelleted through centrifugation at 3,200*g* for 2 minutes.

Forward and reverse primers (Integrated DNA Technologies), for each protein sample were created for downstream reverse transcription and qPCR experiments. Each primer pair was compatible for transcripts produced from the old promoter and T7 Max. For eGFP, the forward primer was 5’-AAGTTCATCTGCACCACC-3’ and the reverse primer was 5’-TTGAAGTCGATGCCCTTC-3’. For the FlAsH peptide, the forward primer was 5’-TATACCGGTATGTGGGACTG-3’ and the reverse primer was 5’-GATGGTGATGATGGTGATGG-3’. For OphA, the forward primer was 5’-ACGACAATGGCAAGTCCA-3’ and the reverse primer was 5’-GGAAATCCGATGCCTCGT-3’.

To prepare the reverse transcription reaction, 2 μL of the DNase-treated sample was mixed with 2 μ L of 10 μ M reverse primer, 4 μ L of 5X Protoscript II Reverse Transcriptase Buffer, 1 μ L of Protoscript II Reverse Transcriptase (200U/μ L, Catalog No. M0368, New England BioLabs Inc.), 2 μ L of 0.1M dithiothreitol (DTT), 1 μ L of 10mM dNTP, 0.2 μ L of RNase Inhibitor (Catalog No. M0314, New England BioLabs Inc.), and 8 μ L of nuclease-free water. The reverse transcription reaction was incubated at 42°C for 1 hour and the reverse transcriptase was inactivated at 65 °C for 20 minutes.

The quantitative PCR reaction mix was prepared by mixing 2 μ L of complementary DNA from the reverse transcription with 2 μL of 10 μ M forward and reverse primers, 11.25 μ L OneTaq Hot Start 2X Master Mix with Standard Buffer (Catalog No. M0484, New England BioLabs Inc.), 1.25 μ L Chai Green Dye 20X (Catalog No. R01200, Chai Bio), and 7.5 μ L of nuclease-free water. The qPCR was completed using Open qPCR (Chai Biotechnologies) with the following thermocycling program: 1 cycle of 30 second denaturation at 95°C, 30 cycles of 15 second denaturation at 95°C, 15 second annealing at 50°C, 1 minute extension at 68°C, and 1 cycle of 5-minute final extension at 68°C. The amplification curves plotted through the Open qPCR software to determine Cq values and averages across 3 replicates of each promoter type were calculated separately.

For experiments involving the kinetic determination of protein expression and transcript comparison, 50 μ L of TXTL reactions with 10 nM DNA templates were incubated at 30°C for 8 hours. Every 2 hours, including at the start of the incubation, 10 μ L samples were removed to measure protein expression and quantify transcription.

### Apta-NASBA reactions

Apta-NASBA reactions were performed as previously described.[31] Primers used for the Apta-NASBA reaction were: Broccoli aptamer coding primer (broccoli is in italics) 5’-*GAGCCCACACTCTACTCGACAGATACGAATATCTGGACCCGACCGTCTC*CAGCGATACATTAAGACGCCTAAAG-3’ classic T7 primer (promoter is in italics) 5’-*TAATACGACTCACTATAG*CGTCAGCATCAGCTACAATTATTCC-3’ T7Max primer (promoter is in italics) 5’-*AATTCTAATACGACTCACTATAGGGA*GACGTCAGCATCAGCTACAATTATTCC-3’

## Acknowledgments

We thank Richard Murray for providing a sample of the GamS protein. We thank Vincent Noireaux for advice on T7 RNAP expression and on using Chi6 inhibitor system. We thank Michael Freeman for the gift of OphA protein from *Omphalotus olearius* Jack-o’-Lantern mushroom.

This work was supported by the NIH award 5R01MH114031, RNA Scaffolds for Cell Specific Multiplexed Neural Observation; NSF award 1840301, RoL:FELS:RAISE: Building and Modeling Synthetic Bacterial Cells; and John Templeton Foundation award 61184, Exploring the Informational Transitions Bridging Inorganic Chemistry and Minimal Life.

## Authors’ contributions

CD, BC, JS, WS, NG, JH, KS and KA performed protein expression experiments. CD, JS, WS and KA wrote the manuscript. All authors analyzed data, edited and approved the manuscript.

## Availability of data and materials

None.

## Consent for publication

All authors agree with publishing this article within The Journal of Biological Engineering

